# Physical and mental health characteristics of adults with subjective cognitive decline: A study of 3,407 people aged 18-81 years from an MTurk-based U.S. national sample

**DOI:** 10.1101/2020.04.07.029553

**Authors:** Ryan Van Patten, Tanya T. Nguyen, Zanjbeel Mahmood, Ellen E. Lee, Rebecca E. Daly, Barton W. Palmer, Tsung-Chin Wu, Xin Tu, Dilip V. Jeste, Elizabeth W. Twamley

## Abstract

Subjective cognitive decline (SCD), or internal feelings of reduced mental capacity, is of increasing interest in the scientific, clinical, and lay community. Much of the extant literature is focused on SCD as a risk factor for Alzheimer’s disease in older adults, while less attention has been paid to non-cognitive health correlates of SCD across adulthood. Consequently, we investigated physical and mental health correlates of SCD in younger, middle-aged, and older adults. We recruited 3,407 U.S. residents through Amazon’s Mechanical Turk, an online labor market. Participants completed a 90-item self-report survey questionnaire assessing sociodemographic characteristics, physical health, sleep, depression, anxiety, loneliness, wisdom, self-efficacy, and happiness. Overall, 493/1930 (25.5%) of younger adults (18-49) and 278/1032 (26.9%) of older adults (50 or older) endorsed the SCD item. Multivariate analysis of variance and follow-up *t*-tests revealed worse physical and mental health characteristics in people endorsing SCD compared to those who did not, with effect sizes primarily in the medium to large range. Additionally, age did not moderate relationships between SCD and physical and mental health. Results suggest that SCD is associated with a diverse set of negative health characteristics such as poor sleep and high body mass index, and lower levels of positive factors including happiness and wisdom. Effect sizes of psychological correlates of SCD were as large as (or larger than) those of physical correlates, indicating that mental health and affective symptoms may be critical to consider when evaluating SCD. Overall, findings from this large, national U.S. sample suggest the presence of relationships between SCD and multiple psychological and perceived health factors; our results also show that SCD may be highly prevalent in both younger and older adults, suggesting that it be assessed across the adult lifespan.

## Introduction

A self-reported decline in cognitive abilities – i.e., subjective cognitive decline (SCD) – is a common complaint in older adults with and without objective cognitive deficits (1). A burgeoning literature in the field of aging research focuses on examining the utility of SCD as an indicator of underlying pathological age-associated cognitive decline years before the onset of the objective, measurable symptoms identifiable in mild cognitive impairment (MCI) and dementia (2). However, whether or not SCD represents an early clinical manifestation of Alzheimer’s disease (AD) pathology remains to be determined. Studies supporting the utility of SCD have found it to be associated with AD neurochemical biomarkers (3). Moreover, in a recent review, Jessen and colleagues (4) concluded that SCD increases risk for pathological cognitive decline on a population level, but that most people with SCD will not convert to MCI and dementia. In contrast, other studies have found inconsistent associations between SCD and objective cognitive functioning in preclinical disease phases (1) and, in some cases, even MCI (5). Moreover, some researchers have reported no relationship between SCD and neuropathological biomarkers of AD (6), and others have found subjective complaints to be an innocuous condition with little risk for future cognitive decline (7–9).

SCD base rates in older adults are high (27-43% in people in their 60s and 70s; 1), evidence for SCD as a risk factor for future cognitive decline is inconclusive (4), and the costs of comprehensive workups for those reporting SCD can be high (10). Moreover, current evidence demonstrates that SCD in the absence of objective cognitive symptoms is associated with worse physical health (11), subjective and objective sleep disturbance (12–14), and psychiatric symptom severity (11,14–16). Negative personality traits (e.g., neuroticism and lower general perceived self-efficacy) have also been linked to SCD in older adults (11,16,17), underscoring the importance of assessing socioemotional health in the context of SCD. Consequently, it is important to investigate non-cognitive correlates of SCD in order to better elucidate the full clinical syndrome and appropriately direct physical and mental healthcare resources.

Relative to older adults, non-cognitive correlates of SCD remain under-investigated in non-clinical samples of younger adults, who are unlikely to experience objective cognitive decline due to neurodegeneration, despite SCD being reported with equal frequency across all ages (18–22). In comparison to older adults who more frequently attribute their SCD to intrinsic, age-associated cognitive decline, younger adults are more likely to attribute SCD to extrinsic, modifiable causes, such as stress, multitasking, and concentration problems (20,21,23). However, the evaluation of SCD in younger adults has been restricted primarily to clinical and medical populations (24–26). The few studies that recruited non-clinical younger and older adults found correlates of SCD to be similar across the two age groups, underscoring the importance of stress, sleep disturbance, and psychiatric symptom severity in SCD across the lifespan (14,19–21,27). However, measures of physical functioning were not consistently and/or comprehensively examined across these studies. Moreover, these studies vary in their recruitment methods and research setting (e.g., memory clinics, surveys explicitly informing participants of the nature of the survey), which may influence prognostically-relevant sample sociodemographic/clinical characteristics (28).

Most of the studies in the exiguous lifespan SCD literature have been conducted in geographically restricted areas (e.g., Korea (27); Paris suburb (21); Portugal (18)), with no prior investigations in a demographically representative and age-diverse U.S. sample. Furthermore, studies within the overall SCD literature have not examined the role of psychological constructs such as wisdom, resilience, and loneliness across the adult lifespan and this information is of essential importance in situating SCD amongst important health-related constructs. As such, the aims of the current study were to comprehensively characterize physical and mental health correlates of SCD across the lifespan in a large, demographically diverse US sample. We recruited participants using Amazon’s Mechanical Turk (AMT), an online labor market allowing for the rapid acquisition of high quality data at low cost (29–33). Based on previously reviewed literature, we hypothesized that, compared to participants who do not endorse SCD (SCD-), those who endorse SCD (SCD+) would report worse physical and mental health.

Furthermore, because of the well-known impact of aging on cognition, we explored the moderating effect of age on the relationship between SCD status and mental/physical health.

## Methods

### Participants

We recruited 3,407 people, aged 18-81, from AMT (see Table 1). Participants completed a 90-item online survey during a five-week period in spring 2019. The description of the survey, visible on AMT read, *“We are looking for people to answer questions about a variety of topics, including age, gender, mood, wisdom, and sleep, among others.”* We described the survey in general terms so as to reduce sampling bias and enhance generalizability. Interested participants consented to the study by selecting a hyperlink, which routed them to the questionnaire, presented via SurveyGizmo.

**Table 1.**
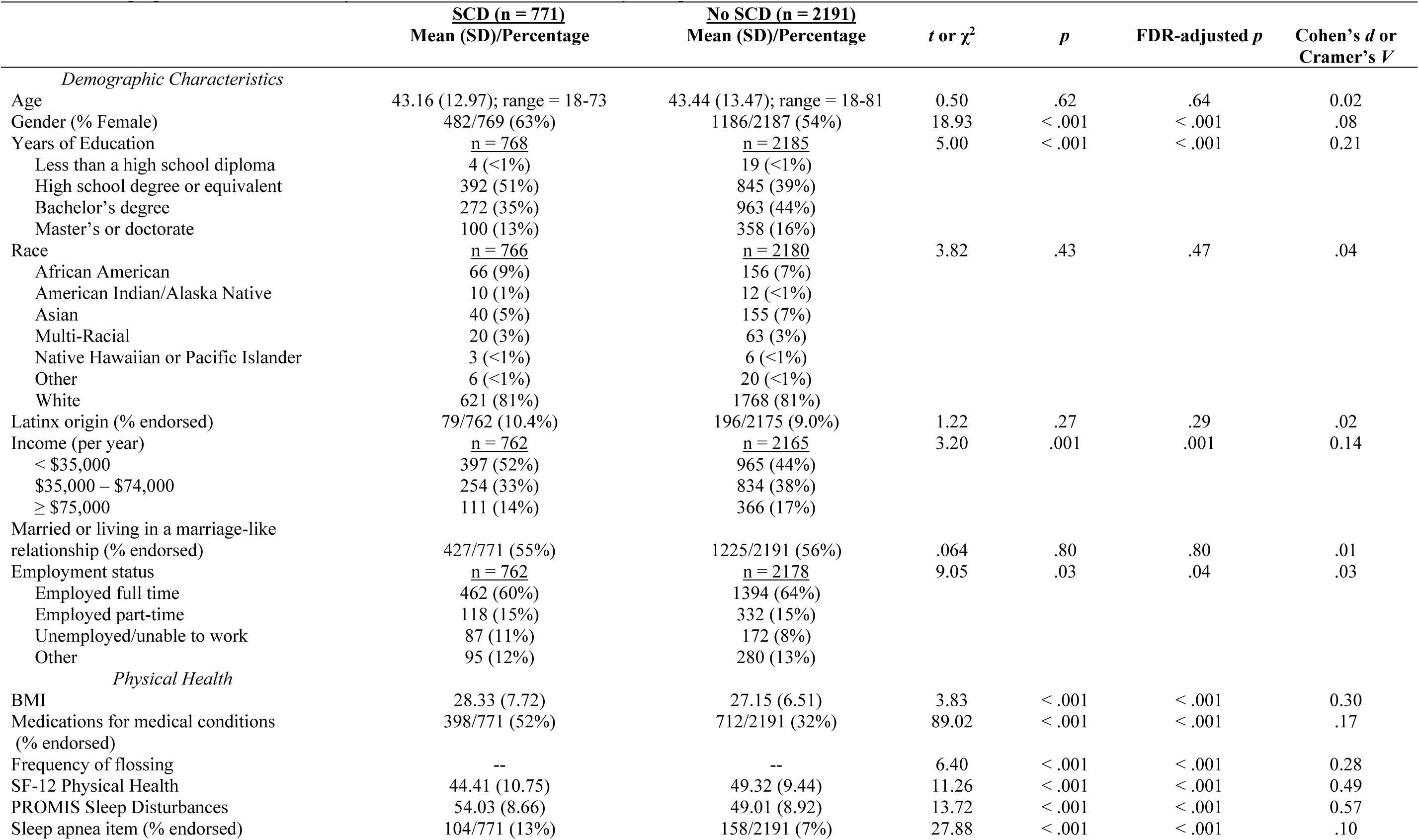
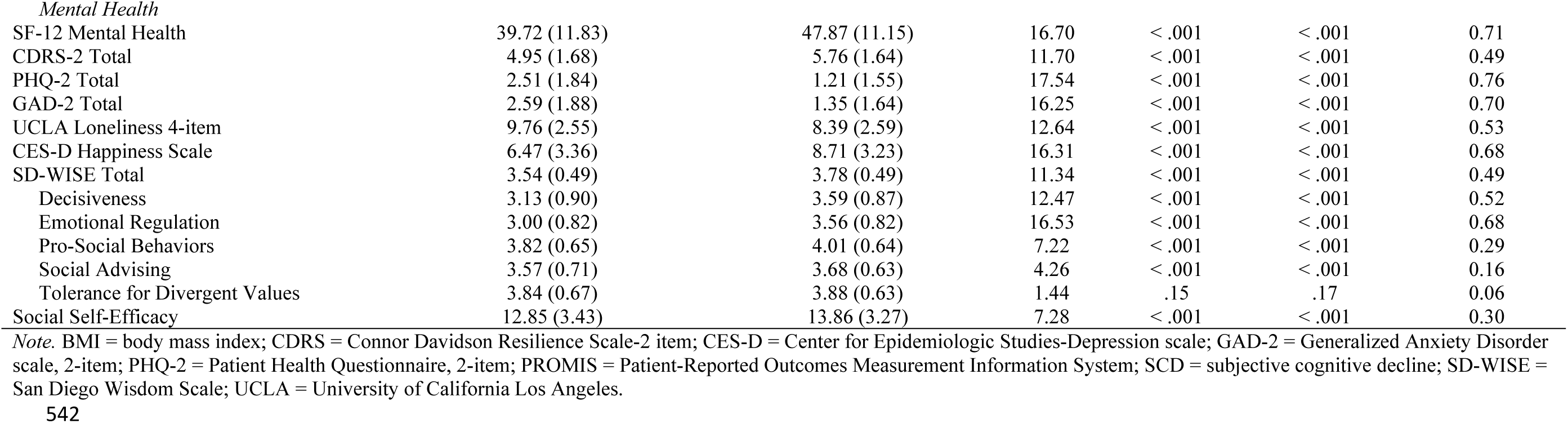
Demographic Characteristics, Physical Health, and Mental Health by Group.

| | **SCD (n = 771)**<br>Mean (SD)/Percentage | **No SCD (n = 2191)**<br>Mean (SD)/Percentage | *t* or $\chi^2$ | *p* | FDR-adjusted *p* | Cohen's *d* or<br>Cramer's *V* |
| --- | --- | --- | --- | --- | --- | --- |
| *Demographic Characteristics* |  |  |  |  |  |  |
| Age | 43.16 (12.97); range = 18-73 | 43.44 (13.47); range = 18-81 | 0.50 | .62 | .64 | 0.02 |
| Gender (% Female) | 482/769 (63%) | 1186/2187 (54%) | 18.93 | < .001 | < .001 | .08 |
| Years of Education | <u>n = 768</u> | <u>n = 2185</u> | 5.00 | < .001 | < .001 | 0.21 |
| Less than a high school diploma | 4 (<1%) | 19 (<1%) |  |  |  |  |
| High school degree or equivalent | 392 (51%) | 845 (39%) |  |  |  |  |
| Bachelor's degree | 272 (35%) | 963 (44%) |  |  |  |  |
| Master's or doctorate | 100 (13%) | 358 (16%) |  |  |  |  |
| Race | <u>n = 766</u> | <u>n = 2180</u> | 3.82 | .43 | .47 | .04 |
| African American | 66 (9%) | 156 (7%) |  |  |  |  |
| American Indian/Alaska Native | 10 (1%) | 12 (<1%) |  |  |  |  |
| Asian | 40 (5%) | 155 (7%) |  |  |  |  |
| Multi-Racial | 20 (3%) | 63 (3%) |  |  |  |  |
| Native Hawaiian or Pacific Islander | 3 (<1%) | 6 (<1%) |  |  |  |  |
| Other | 6 (<1%) | 20 (<1%) |  |  |  |  |
| White | 621 (81%) | 1768 (81%) |  |  |  |  |
| Latinx origin (% endorsed) | 79/762 (10.4%) | 196/2175 (9.0%) | 1.22 | .27 | .29 | .02 |
| Income (per year) | <u>n = 762</u> | <u>n = 2165</u> | 3.20 | .001 | .001 | 0.14 |
| < \$35,000 | 397 (52%) | 965 (44%) | | | | |
| \$35,000 – \$74,000 | 254 (33%) | 834 (38%) | | | | |
| ≥ \$75,000 | 111 (14%) | 366 (17%) | | | | |
| Married or living in a marriage-like<br>relationship (% endorsed) | 427/771 (55%) | 1225/2191 (56%) | .064 | .80 | .80 | .01 |
| Employment status | <u>n = 762</u> | <u>n = 2178</u> | 9.05 | .03 | .04 | .03 |
| Employed full time | 462 (60%) | 1394 (64%) |  |  |  |  |
| Employed part-time | 118 (15%) | 332 (15%) |  |  |  |  |
| Unemployed/unable to work | 87 (11%) | 172 (8%) |  |  |  |  |
| Other | 95 (12%) | 280 (13%) |  |  |  |  |
| *Physical Health* |  |  |  |  |  |  |
| BMI | 28.33 (7.72) | 27.15 (6.51) | 3.83 | < .001 | < .001 | 0.30 |
| Medications for medical conditions<br>(% endorsed) | 398/771 (52%) | 712/2191 (32%) | 89.02 | < .001 | < .001 | .17 |
| Frequency of flossing | -- | -- | 6.40 | < .001 | < .001 | 0.28 |
| SF-12 Physical Health | 44.41 (10.75) | 49.32 (9.44) | 11.26 | < .001 | < .001 | 0.49 |
| PROMIS Sleep Disturbances | 54.03 (8.66) | 49.01 (8.92) | 13.72 | < .001 | < .001 | 0.57 |
| Sleep apnea item (% endorsed) | 104/771 (13%) | 158/2191 (7%) | 27.88 | < .001 | < .001 | .10 |
| *Mental Health* |  |  |  |  |  |  |
| SF-12 Mental Health | 39.72 (11.83) | 47.87 (11.15) | 16.70 | < .001 | < .001 | 0.71 |
| CDRS-2 Total | 4.95 (1.68) | 5.76 (1.64) | 11.70 | < .001 | < .001 | 0.49 |
| PHQ-2 Total | 2.51 (1.84) | 1.21 (1.55) | 17.54 | < .001 | < .001 | 0.76 |
| GAD-2 Total | 2.59 (1.88) | 1.35 (1.64) | 16.25 | < .001 | < .001 | 0.70 |
| UCLA Loneliness 4-item | 9.76 (2.55) | 8.39 (2.59) | 12.64 | < .001 | < .001 | 0.53 |
| CES-D Happiness Scale | 6.47 (3.36) | 8.71 (3.23) | 16.31 | < .001 | < .001 | 0.68 |
| SD-WISE Total | 3.54 (0.49) | 3.78 (0.49) | 11.34 | < .001 | < .001 | 0.49 |
| Decisiveness | 3.13 (0.90) | 3.59 (0.87) | 12.47 | < .001 | < .001 | 0.52 |
| Emotional Regulation | 3.00 (0.82) | 3.56 (0.82) | 16.53 | < .001 | < .001 | 0.68 |
| Pro-Social Behaviors | 3.82 (0.65) | 4.01 (0.64) | 7.22 | < .001 | < .001 | 0.29 |
| Social Advising | 3.57 (0.71) | 3.68 (0.63) | 4.26 | < .001 | < .001 | 0.16 |
| Tolerance for Divergent Values | 3.84 (0.67) | 3.88 (0.63) | 1.44 | .15 | .17 | 0.06 |
| Social Self-Efficacy | 12.85 (3.43) | 13.86 (3.27) | 7.28 | < .001 | < .001 | 0.30 |

*Note.* BMI = body mass index; CDRS = Connor Davidson Resilience Scale-2 item; CES-D = Center for Epidemiologic Studies-Depression scale; GAD-2 = Generalized Anxiety Disorder scale, 2-item; PHQ-2 = Patient Health Questionnaire, 2-item; PROMIS = Patient-Reported Outcomes Measurement Information System; SCD = subjective cognitive decline; SD-WISE = San Diego Wisdom Scale; UCLA = University of California Los Angeles.

We paid each worker $1.00 for survey completion. Inclusion criteria were the following: 1) ≥18 years old, 2) English-speaking, 3) residing in the U.S., and 4) a Human Intelligence Task Approval rate >90% (32). We initially recruited 2,289 participants and found that the age distribution was skewed toward younger adults. In order to balance the sample with respect to age, we added 250 more participants aged 35-45, 500 participants aged 45-55, and 368 participants aged 55+, leading to the initial sample size of 3,407.

Although AMT workers provide high quality data overall (29,31,33–38), a subset may be inattentive or may provide invalid data for other reasons. In order to address this issue, we excluded participants who provided impossible or highly improbable answers to survey items. Specifically, we excluded participants who, 1) completed the survey in <270 seconds (n=104), 2) reported values for height and weight leading to a body mass index (BMI)<16 (n=165), 3) reported fewer total close friends than the number of close friends seen at least once per month (n=252), 4) reported their height at <3 feet or >7 feet (n=42), 5) reported living with ≥20 people in their household (n=12), and/or 6) reported owning ≥40 pets (n=3). Overall, 336 participants provided one invalid response, 86 participants provided two invalid responses, 22 participants provided three invalid responses, and 1 participant provided four invalid responses. Applying these exclusion criteria resulted in 445 (13.1%) participants being excluded, leaving a final sample of 2,962 participants for analysis.

This project, including a request for a waiver of documented consent, was reviewed through the UC San Diego Human Research Protections Program by an IRB Chair and/or the IRB Chair’s designee and certified as exempt from IRB review under 45 CFR 46.104(d), Category 2.

### Materials

Within the 90-item survey, we measured SCD with a single item (*“Have you noticed a decline in your memory and thinking that is worrisome to you? [Yes/No]”*). We attempted to minimize the length of the survey as much as possible; consequently, we selected empirically-supported abbreviated versions of all measures with the exception of the San Diego Wisdom Scale (SD-WISE), which does not have a short form. We measured multiple sociodemographic characteristics including age, sex, education, race, annual income, marital status, and employment status. To assess physical health, we administered one item inquiring about frequency of flossing (once per week, 2-3 times per week, 4-6 times per week, or daily; 39), two items measuring height and weight to calculate body mass index (BMI), one item asking whether or not any medications are taken for medical conditions, the 12-item Medical Outcomes Survey-Short Form (assessing physical and mental health related quality of well-being; 39), the 4-item Patient-Reported Outcomes Measurement Information System (PROMIS) Sleep Disturbance-short form (41), and the 1-item PROMIS sleep apnea question (41). We measured depression with the 2-item Patient Health Questionnaire (PHQ-2; 42) and anxiety with the 2-item version Generalized Anxiety Disorder scale (43). We measured loneliness with the 4-item version of the UCLA Loneliness Scale (44), using the anchors from the third edition of the UCLA scale, *never, rarely, sometimes*, and *always*, rather than those of Russel et al. (44), *never, rarely, sometimes*, and *often*. Measures of positive psychological factors included the 24-item SD-WISE (45), the 2-item Connor-Davidson Resilience Scale (46), and the 4-item Happiness Factor from the Center for Epidemiologic Studies-Depression scale (47). The SD-WISE includes the following subscales: Decisiveness, Emotional Regulation, Pro-Social Behaviors, Social Advising, and Tolerance for Divergent Values. We assessed social self-efficacy using four items, with minor wording modifications, from the Social Self-Efficacy Scale (48,49) that was originally developed for use with adolescents. These four items were selected for age-appropriateness and included: (1) “*How well can you become friends with other people?*,” (2) “*How well can you have a chat with an unfamiliar person?*,” (3) “*How well can you tell other people that they are doing something you don’t like?*,” and (4) “*How well can you succeed in preventing quarrels with other people?*”.

### Statistical Analyses

We analyzed the data using SPSS 26.0. We first examined distributional characteristics of all continuous variables. For those that were highly skewed, we used appropriate non-parametric tests. Next, we tested our hypothesis with SCD group as the between-subjects predictor (“independent”) variable and physical and mental health scores as outcome (“dependent”) variables Casual inference cannot be established with this cross-sectional data, but our main focus was to test whether SCD status predicted physical and mental health levels. We ran an omnibus multivariate analysis of variance (MANOVA), followed by independent samples *t*-tests for each continuous variable; due to low missing data rates (<4% for all variables and <1% for all variables except for frequency of flossing), we used the classic MANOVA procedure rather than the generalized estimating equations procedure (50). For the two categorical outcome variables (presence or absence of medications and sleep apnea), we conducted χ^2^ tests. With regard to the exploratory analysis, we conducted 2-group (SCD+ versus SCD-) X 2-age cohort (older: 50+ versus younger: 18-49) ANOVAs on the physical and mental health variables listed in Table 1 in order to examine the possible moderating effect of age. We dichotomized age into two groups in order to contrast younger adults with older adults, given that the majority of the current SCD literature exists in aging populations. Although some researchers define older adults beginning in the 60s, we were interested in “young-old” adults, and our sample of “old-old” adults was small, likely due to our use of an internet-based data collection platform (AMT).

We report appropriate effect sizes for all statistical tests – partial η^2^ for the MANOVA, Cohen’s *d* for *t* tests, and Cramer’s *V* for χ^2^ tests. We used the False Discovery Rate to control for Type I error, with alpha set at *p*<.05. The False Discovery Rate predicts and controls individual false positive results, while simultaneously maintaining a high level of statistical power relative to familywise error rate methods such as the Bonferroni correction (51). All statistical tests were two-tailed.

## Results

Overall, 493/1930 (25.5%) of younger adults (18-49) and 278/1032 (26.9%) of older adults (50 or older) endorsed the SCD item, χ^2^(1)=.68, *p*=.41. For continuous variables with non-normal distributions, results from non-parametric tests (Mann Whitney *U*s) mirrored those from parametric statistics. For ease of interpretation, we present the parametric results for all continuous variables. Of the demographic variables, sex, education, employment status, and annual income differed significantly across the two groups; however, when we added these variables to the models as covariates, results did not differ. For ease of interpretation, we provide unadjusted parameters.

With respect to our hypothesis that the SCD+ group would report worse physical and mental health compared to the SCD-group, the MANOVA was statistically significant, *F*(11, 2937)=46.47, *p*<.001, λ=0.85, 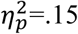. In univariate analyses, all physical and mental health variables differed in the hypothesized direction, with the exception of SD-WISE Tolerance for Diverging Values subscale. Specifically, compared to the SCD-group, the SCD+ group reported higher BMI, greater rate of taking medications for medical conditions, less frequent flossing, worse overall self-reported physical health, higher rates of self-reported sleep disturbance and sleep apnea, worse overall mental health, higher rates of depression, anxiety, and loneliness, and lower scores on scales for resilience, happiness, wisdom, and self-efficacy (see Table 1). Moreover Cohen’s *d* effect sizes were primarily in the medium (0.50) to large (0.80) range. When we split the sample by age (18-49 and 50+) for the exploratory analysis, interaction terms for the 2-SCD group X 2-Age group ANOVAs were all non-significant, suggesting that age did not moderate the relationship between SCD and physical or mental health.

## Discussion

Most research efforts to investigate SCD have been focused on understanding its relationship to objective cognitive decline (52,53) and its importance as an early risk factor for MCI and dementia in older adults (4,54). Although the literature on SCD as a marker of early cognitive decline in older adults is growing rapidly, much less is known about SCD as a general construct, especially its physical and mental health correlates in non-clinical populations across the adult lifespan. The present study evaluated self-reported physical and psychological correlates of SCD in a large survey sample of adults aged 18-81. As hypothesized, both younger and older adults who endorsed SCD exhibited worse self-reported physical health symptoms and psychological traits/states compared to those who did not endorse SCD. Compared to SCD-participants, SCD+ participants had higher mean BMI, were more likely to take medications for medical conditions, were more likely to have sleep apnea or other sleep disturbances, and were less likely to floss. They also reported worse physical well-being, worse mental well-being, higher depression and anxiety symptoms, greater loneliness, and lower levels of resilience, happiness, wisdom, and self-efficacy. Age did not moderate the relationship between SCD and either physical or psychological functioning. Overall, our findings are consistent with previous literature suggesting that SCD is associated with worse physical health (11), subjective and objective sleep disturbance (12–14), and psychiatric symptom severity (11,14–16), and that SCD correlates are similar across the lifespan in both younger and older adults (19,21,27). To our knowledge, this study is the first large-scale investigation of SCD rates to include a non-clinical sample of younger adults in the U.S. Notably, the prevalence of SCD did not differ between younger and older adults, which was unexpected. Previous studies have reported similar rates of SCD in younger adults (approximately 25-29%) but higher rates in older adults (20,27). A possible explanation for the lack of difference in SCD between younger and older adults is that our older adult sample recruited through AMT may be different than those in other clinical studies. Nevertheless, other studies have found SCD to be as frequent, though qualitatively different, in young adults compared to older adults (18,21).

Depression and anxiety symptoms demonstrated the strongest relationships to SCD, with moderate-to-large (Cohen’s *d* ≥ 0.70) effect sizes. This finding is consistent with earlier studies, which suggest that the relationship between SCD and symptoms of depression and anxiety is complex. Depression moderates the relationship between SCD and objective cognitive impairment (14,55). Moreover, although depression and anxiety are closely related and highly comorbid, they are associated with different risk factors; pure anxiety tends to be associated with a wide range of stress-related factors, none of which are associated with pure depression (56). One factor common to both is personal mastery, or perceived behavioral control, which may be a cognitive psychological marker of trait vulnerability for both depression and anxiety (56). Depression interacts with personal mastery and general self-efficacy such that the association between depressive symptoms and SCD may be stronger in participants with higher feelings of perceived mastery and social self-efficacy (11). Our data revealed that individuals who endorsed SCD exhibited lower levels of self-efficacy. Memory complaints may reflect a general state of diminished psychological or mental well-being, which was also observed in this study.

We also observed strong associations between SCD and negative/positive psychological factors, including loneliness, resilience, happiness, and wisdom (Cohen’s *d* ≥ 0.30). In each case, SCD was related to higher levels of negative and lower levels of positive psychological factors, and effect sizes of psychological correlates were as large as (or larger) than those of physical correlates, suggesting that psychological features may be associated with SCD as much as physical functioning. These results represent a unique contribution to the current SCD literature and have important clinical implications. The associations between SCD and negative/positive psychological factors point to possible areas of intervention. For example, increasing one’s subjective cognitive experience may improve resilience and happiness and decrease levels of loneliness; conversely, interventions aimed at improving psychological factors may improve one’s subjective cognitive experience.

Wisdom is a complex, multidimensional personality trait that is comprised of several specific components, including pro-social behaviors such as empathy and compassion, emotional regulation, self-reflection or insight, acceptance of divergent values, decisiveness, and social advising (57–59). Although it is often conflated with intelligence, wisdom encompasses cognitive, affective, and reflective dimensions (60). Among the wisdom subscales of the SD-WISE, the cognitive (decisiveness) and affective (emotional regulation) components of wisdom were the strongest correlates of SCD. The decisiveness component entails the cognitive abilities and dispositional qualities related to making decisions. The emotional regulation component pertains to the ability to maintain emotional homeostasis. Although the latter can be reflective of psychological distress, one of the items (e.g., *I cannot filter my negative emotions*) also involves an aspect impulse control related to frontal executive functions, specifically response inhibitory (57). Thus, it is not surprising that individuals with SCD would have lower decisiveness and emotional regulation. At the same time, it is worth stress that positive traits are potentially modifiable. There is growing literature on interventions designed to enhance levels of positive traits such as resilience and components of wisdom including emotional regulation (61,62).

In addition, individuals with SCD reported greater loneliness, which has been previously identified as a major risk factor for adverse mental and physical health outcomes, including cognitive decline and dementia (63–66). (Please see our companion paper, Nguyen et al. (67), for description of detailed analyses of loneliness and associated factors within this MTurk sample.)

With regard to physical functioning, SCD had the strongest relationship with self-reported sleep disturbances and overall physical well-being, consistent with previous literature (27). Disrupted sleep can contribute to both subjective and objective experiences of cognitive impairment (12,68). Similarly, presence of sleep disorders – namely, obstructive sleep apnea (OSA) – has been associated with SCD. Although cognitive deficits have been well documented in patients with OSA, the relationship between SCD and objective impairment in OSA remains unclear (69). SCD in combination with subjective sleep disturbance and OSA may represent early prodromal signs for developing MCI or dementia (70).

The present study has notable strengths. It includes a large sample of nearly 3,000 adults across the adult lifespan with sociodemographic diversity in terms of gender, race, and socioeconomic status. Utilization of the AMT online crowdsourcing marketplace allowed for access to thousands of research participants from demographically diverse backgrounds from around the US, without geographic restrictions (29). Recruitment through internet samples may reduce biases from traditional samples (71) and better approximate US census data (72–76). Moreover, we took the precaution of using general terms to describe the survey so as to reduce sampling bias and enhance generalizability. Although the unsupervised nature of data collection potentially reduces reliability and validity, many studies have shown that AMT data quality is equivalent to that acquired in controlled settings (29,31–33,38,76) and excluded participants who provided impossible or highly implausible responses to survey items to ensure validity of results. Furthermore, our study included a comprehensive assessment of physical and mental health factors, including positive and negative psychological traits/states, which, to our knowledge, have never been simultaneously investigated in the context of SCD. Overall, our findings provide a more comprehensive understanding of the physical and psychological characteristics, above and beyond psychopathology, associated with SCD.

Nevertheless, this investigation also had several limitations. The presence of SCD was determined using a single yes-no question, rather than a more detailed method of inquiry or standardized measure, which restricted our ability to assess SCD severity and capture complaints in specific cognitive domains. Many self-report measures have been used to investigate SCD (77), but there is no established gold standard method of assessment (78). Moreover, in our experience, this mode of assessment is more pragmatic and consistent with typical clinical practice, and most individuals with impairments in other cognitive domains (e.g., attention/concentration, language, executive function) often perceive these problems as memory difficulties. Due to restrictions of the AMT platform, all data are self-report, which has well known limitations due to recall and response bias (79,80). Although the assessment of subjective cognitive decline requires self-report by definition, this represents a limitation with regard to reports of physical health and functioning. Relatedly, we did not administer performance-based cognitive tests to determine objective cognitive impairment. Finally, the cross-sectional design limits our ability to draw causal inferences regarding SCD and its correlates, and future prospective longitudinal studies are needed to clarify these relationships.

## Conclusions

Notwithstanding these limitations, the current study contributes to important research aimed at better understanding the non-cognitive aspects of SCD. Our findings help to characterize the wide range of physical and psychological correlates of SCD. Notably, although definitive causal conclusions are limited by reliance on subjective self-reports, the effect sizes of psychological correlates of SCD were as large as (or larger) than those of physical correlates, indicating that mental health and psychological features are critical to consider when evaluating SCD.

## Acknowledgements

The authors thank all the study participants for their contributions to this work.

